# Neuronal *SNCA* transcription during Lewy body formation

**DOI:** 10.1101/2023.08.19.553427

**Authors:** Tomoya Kon, Shelley L. Forrest, Seojin Lee, Ivan Martinez-Valbuena, Jun Li, Nasna Nassir, Mohammed J. Uddin, Anthony E. Lang, Gabor G. Kovacs

**Author notes:** **Correspondence:** Gabor G. Kovacs, MD, PhD, FRCPC. Tanz Centre for Research in Neurodegenerative Disease, University of Toronto, Toronto, ON, Canada 60 Leonard Ave., Rm 6KD414, Tanz CRND, Krembil Discovery Tower, University of Toronto, Toronto, Ontario M5T 0S8, Canada.

## Abstract

**Background:** Misfolded α-synuclein (α-syn) is believed to contribute to neurodegeneration in Lewy body disease (LBD) based on considerable evidence including a gene-dosage effect observed in relation to point mutations and multiplication of *SNCA* in familial Parkinson’s disease. A contradictory concept proposes early loss of the physiological α-syn as the major driver of neurodegeneration. There is a paucity of data on *SNCA* transcripts in various α-syn immunoreactive cytopathologies.

**Methods:** *SNCA* transcripts in neurons without and with various α-syn immunoreactive cytopathologies in the substantia nigra and amygdala in LBD (n = 5) were evaluated using RNAscope combined with immunofluorescence for disease-associated α-syn. Single-nucleus RNA sequencing was performed to elucidate cell-type specific *SNCA* expression in non-diseased frontal cortex (n = 3).

**Results:** *SNCA* transcripts in neurons with punctate α-syn immunoreactivity were preserved both in the substantia nigra and amygdala but were reduced in neurons with compact α-syn inclusions. Only single *SNCA* transcripts were detected in astrocytes with or without α-syn immunoreactivity in the amygdala. Single-nucleus RNA sequencing revealed that excitatory and inhibitory neurons, oligodendrocyte progenitor cells, oligodendrocytes, and homeostatic microglia expressed *SNCA* transcripts, while expression was largely absent in astrocytes and microglia.

**Conclusions:** The preserved cellular *SNCA* expression in the more abundant non-Lewy body type α-syn cytopathologies provides a pool for local protein production that can aggregate and serve as a seed for misfolded α-syn. Successful segregation of disease-associated α-syn is associated with the exhaustion of *SNCA* production in the terminal cytopathology, the Lewy body. Our observations support a therapeutic strategy incorporating a finely tuned dual approach targeting the elimination of misfolded α-syn along with the reduction of the *SNCA* transcription to avoid feeding of pathological α-syn seeding.

## Introduction

Lewy body disease (LBD) is characterized by the presence of Lewy bodies (LBs) composed of α-synuclein (α-syn) [1, 2]. α-Syn is a neuronal cytoplasmic and pre-synaptic protein encoded by *SNCA* gene, which was originally described in the pre-synaptic terminal and nucleus of neurons from Torpedo californica [3]. A conformational change, termed also as misfolding, gives rise to the emergence of an aggregation nucleus composed of α-syn, commonly referred to as a seed [2, 4]. This seed possesses the capacity to actively engage endogenous monomeric α-syn molecules and instigate their aggregation process [5-7]. The application of α-syn immunohistochemistry allowed the detection of non-LB type cytopathologies that are more abundant than classical LBs. In addition, it suggested that the maturation process of classical LB formation encompasses sequential stages [2, 8-10]. Importantly, cell culture and animal studies [11-14] also showed a similar process. Accordingly, the initial phase of this process is characterized by the presence of punctate (or diffuse) α-syn immunoreactivity (IR) in the cytoplasm, which shows negative IR for both ubiquitin and p62 antibodies. Subsequently, irregular-shaped compact inclusions considered to be analogous to the pale bodies observed in the HE-staining, which exhibit IR against ubiquitin and p62 antibodies, are generated. Ultimately, the progression leads to the formation of fully mature classical LBs characterized by a central round core surrounded by a halo [1, 2, 8, 9, 15].

Rare point mutations and multiplications (duplication and triplication) of *SNCA* lead to familial Parkinson’s disease leading to the hypothesis of a gene-dosage effect [16-19]. Many studies support the notion that accumulation of misfolded α-syn drives disease pathogenesis (‘proteinopathy’) [1, 2, 4–9, 11, 15, 20, 21]. Therefore, α-syn is currently a major therapeutic target for synucleinopathies, for example by eliminating pathological aggregates by monoclonal α-syn antibodies, by inhibiting α-syn aggregation, or by stabilizing α-syn monomers, or by directly targeting *SNCA* gene expression with miRNA and antisense oligonucleotide therapies, amongst others [22-24]. However, it has been argued that the lack of physiological protein (referred to as ‘proteinopenia’) is a major contributor to the neurodegeneration in LBD [25], supported in part by the absence of notable impact in recent clinical trials involving monoclonal α-syn antibodies [22, 23]. These two divergent concepts dictate completely opposing strategies for disease-modifying treatment, specifically the eradication of disease-associated α-syn or the early replacement of normal physiological α- syn.

*SNCA* expression in the substantia nigra (SN) in LBD has been reported to be both up-and down-regulated compared with controls [26-35]. However, these studies did not compare *SNCA* transcription with or without α-syn pathology because tissue digestion methods were used. Elucidation of *SNCA* transcriptional regulation during LB formation process is crucial for understanding the pathogenesis and progression of LBD, and for guiding the strategy for molecular therapy. Here, we report the process of cellular *SNCA* transcription during LB formation using RNAscope combined with immunofluorescence for disease-associated α-syn, complemented by single-nucleus RNA sequencing (snRNA-seq) to map the cell population showing *SNCA* transcripts.

## Methods

### Cases and tissue preparation

Formalin-fixed paraffin-embedded 4-μm thick sections were investigated. For RNAscope, sections were trimmed to approximately 1.5 cm x 1.5 cm in size from the substantia nigra (SN, n = 5 in LBD cases, n = 2 in controls; patients without neurodegenerative pathology), amygdala (n = 3 in LBD, n = 2 in control cases). To compare nuclear and cytoplasmic *SNCA* transcripts, pons (n = 2 in control cases) was also investigated. For immunohistochemistry, midbrain and basal ganglia were analyzed (n = 5 each in cases of LBD and controls). In addition, flash-frozen frontal cortex from 3 control cases stored at –80℃ were also used for snRNA-seq analysis. A total of 13 (5 LBD and 8 control) cases were included in this study (**Table 1**). Non-diseased control samples were used only to demonstrate the presence of *SNCA* transcripts in cells and to compare nuclear and cytoplasmic transcripts. All cases had a routine neuropathological assessment based on the current consensus criteria including Braak LBD stage [20], Lewy pathology consensus criteria [36], and Alzheimer’s disease neuropathological change (ADNC) [37].

**Table 1.** Case demographics, regions, antibodies, and probes used in the study.

| Case | Age | Sex | PMI,<br>hours | Diagnosis | Braak LBD<br>stage<sup>25</sup> | Regions and probes<br>used for RNAscope | Region and antibodies<br>used for IHC | Region used for<br>single-nucleus RNA<br>sequencing |
| --- | --- | --- | --- | --- | --- | --- | --- | --- |
| 1 | 73 | F | n.a. | LBD | 4 | SN (*SNCA*, *RBFOX3*),<br>amygdala (*SNCA*,<br>*RBFOX3*, *ALDH1L1*) | SN and basal ganglia<br>(SYN-1, 5G4) | - |
| 2 | 62 | F | 15 | LBD | 5 | SN (*SNCA*, *RBFOX3*),<br>amygdala (*SNCA*,<br>*RBFOX3*, *ALDH1L1*) | SN and basal ganglia<br>(SYN-1, 5G4) | - |
| 3 | 81 | M | < 24 | LBD | 5 | SN (*SNCA*, *RBFOX3*),<br>amygdala (*SNCA*,<br>*RBFOX3*, *ALDH1L1*) | SN and basal ganglia<br>(SYN-1, 5G4) | - |
| 4 | 62 | M | 4 | LBD | 4 | SN (*SNCA*, *RBFOX3*) | SN and basal ganglia<br>(SYN-1, 5G4) | - |
| 5 | 85 | M | n.a. | LBD | 4 | SN (*SNCA*, *RBFOX3*) | SN and basal ganglia<br>(SYN-1, 5G4) | - |
| 6 | 52 | F | 38 | Control | None | SN (*SNCA*, *RBFOX3*),<br>amygdala (*SNCA*,<br>*RBFOX3*, *ALDH1L1*),<br>pons (*SNCA*, *Olig2*) | SN and basal ganglia<br>(SYN-1, 5G4) | - |
| 7 | 64 | F | n.a. | Control | None | SN (*SNCA*, *RBFOX3*),<br>amygdala (*SNCA*,<br>*RBFOX3*, *ALDH1L1*),<br>pons (*SNCA*, *Olig2*) | SN and basal ganglia<br>(SYN-1, 5G4) | - |
| 8 | 66 | M | 12 | Control | None | - | SN and basal ganglia<br>(SYN-1, 5G4) | - |
| 9 | 66 | M | 20 | Control | None | - | SN and basal ganglia<br>(SYN-1, 5G4) | - |
| 10 | 75 | M | 30 | Control | None | - | SN and basal ganglia<br>(SYN-1, 5G4) | - |
| 11 | 74 | M | 12.5 | Control | None | - | - | Frontal cortex |
| 12 | 74 | M | 6.3 | Control | None | - | - | Frontal cortex |
| 13 | 77 | M | 4 | Control | None | - | - | Frontal cortex |

Abbreviations: IHC, immunohistochemistry; LBD, Lewy body disease; LPC, Lewy pathology consensus criteria; n.a., not available;
PMI, postmortem interval; SN, substantia nigra

### RNAscope with immunofluorescence

RNAscope assay with immunofluorescence was performed as previously reported [38]. Briefly, endogenous peroxidase was quenched by RNAscope Hydrogen Peroxide Solution (ACD) for 10 minutes, and sections were pretreated with RNAscope Target Retrieval reagent (ACD) for 30 minutes at 99 ℃ before applying RNAscope Protease Plus (ACD) for 30 minutes at 40 ℃. Sections were incubated with probe mixtures for 2 hours at 40 °C. C1 probe designates *SNCA* (ACD, Cat no. 421311, targeted regions 291 – 3084 bp, accession number NM_001146054.1) and was detected with Opal 570 fluorophore (1:750). C2 probe labeled *Olig2* for an oligodendrocyte maker (ACD, Cat no. 424191-C2, target regions 929 – 2502 bp, accession number NM_005806.3), C3 probe was assigned for *ALDH1L1* for a marker of astrocytes (ACD, Cat no. 438881-C3, targeted regions 1999 – 2982 bp, accession number NM_001270364.1), and C4 probe was set for *RBFOX3* for a neuronal marker (ACD, Cat no. 415591-C4, targeted regions 720 – 2217 bp, accession number NM_001082575.2). Cell-type-specific probes were detected with the Opal 690 fluorophore (1:750). After probe hybridization, the slides were washed and hybridized, and developed with an RNAscope Multiplex Fluorescent V2 Assay kit. The slides from the midbrain and amygdala were further processed for immunofluorescence using phosphorylated α-syn antibody (clone #64, 1:5000, Wako, Osaka, Japan) for 1 hour incubation at room temperature (RT). To evaluate astrocytic α-syn pathology that is undetectable using phosphorylated α-syn antibodies, for selected amygdala sections, we used the 5G4 α-syn antibody (1:100) for 1 hour incubation at RT [21, 39]. Eighty-percent formic acid for 5 minutes was added before the process of RNAscope protease plus for 5G4 α-syn antibody. After washes, the sections were incubated with Alexa 488-conjugated donkey anti-mouse antibody (1:500, Invitrogen/ThermoFisher) for 1 hour at RT and mounted with 4′,6-diamidino-2-phenylindole (DAPI) with mounting medium. The regions, probes, and antibodies used in the study were summarized in **Table 1**.

### Acquisition of images

Acquisition of images was performed using Nikon C2Si+ confocal on a Nikon Ti2-E inverted microscope equipped with a 40X objective lens for a single stack (NA: 0.95). Appropriate filter settings were used as follows; DAPI, excitation 405 nm, emission 400—720 nm; Alexa 488, excitation 488 nm, emission 430—500 nm; Opal 590, excitation 561 nm, emission 520—600 nm; Opal 690, excitation 640 nm, emission 620—720 nm. Single-stack images were captured using the NIS-Elements AR software (version 5.30.04). Only a few selected sections were also captured with a 100X objective lens for the acquisition of a z-stack of slices, where the inter-slice distance was 0.1 μm. Subsequently, a three-dimensional image was generated utilizing the NIS-Elements AR software in these selected sections. The parameters for taking pictures were standardized and maintained throughout the entire experiment.

### Morphometry

We identified the cells with cell-specific markers for neurons, oligodendrocytes, and astrocytes using cell-specific probes. The region of interest (ROI) for the cell body was manually delineated using the cell-specific markers and DAPI signal positivity that allowed the identification of the contour of the neuronal cytoplasm (**Fig. S1)**. First, the cell-specific marker and DAPI channels were selected and merged. Subsequently, the Lookup Table (LUT) strength was increased sufficiently to enhance the visibility of autofluorescence from neuromelanin and/or lipofuscin pigments. The borders of the cell body were then traced based on these signals (**Fig. S1B)**. Finally, the LUT parameters were restored to normal settings, and the *SNCA* transcript channel is selected (**Fig. S1C**). Finally, the NIS-Elements software captured the area positive for *SNCA* transcripts above the threshold within the ROI (**Fig. S1D**). Following our previous report [38], we established a capture threshold for *SNCA* fluorescence intensity that exceeded the autofluorescence level (**Fig. S1D**). This criterion ensured independence from the potential impact of autofluorescence during analysis. *SNCA* transcripts were identified either as individual entities or in small confluent clusters (**Fig. 1, 2, S1**), making it difficult to accurately count individual transcripts. Consequently, in this study, we relied on the area density values of *SNCA* transcripts [38]. *SNCA* area density was determined by dividing the area occupied by *SNCA* transcripts by the annotated cell body area (ROI) and was displayed as the percentage. The individual measurement of *SNCA* area density was pooled into distinct morphological groups of inclusion types. The parameters used for image analysis were standardized and consistently applied throughout the entire experiment.

**Fig. 1.**
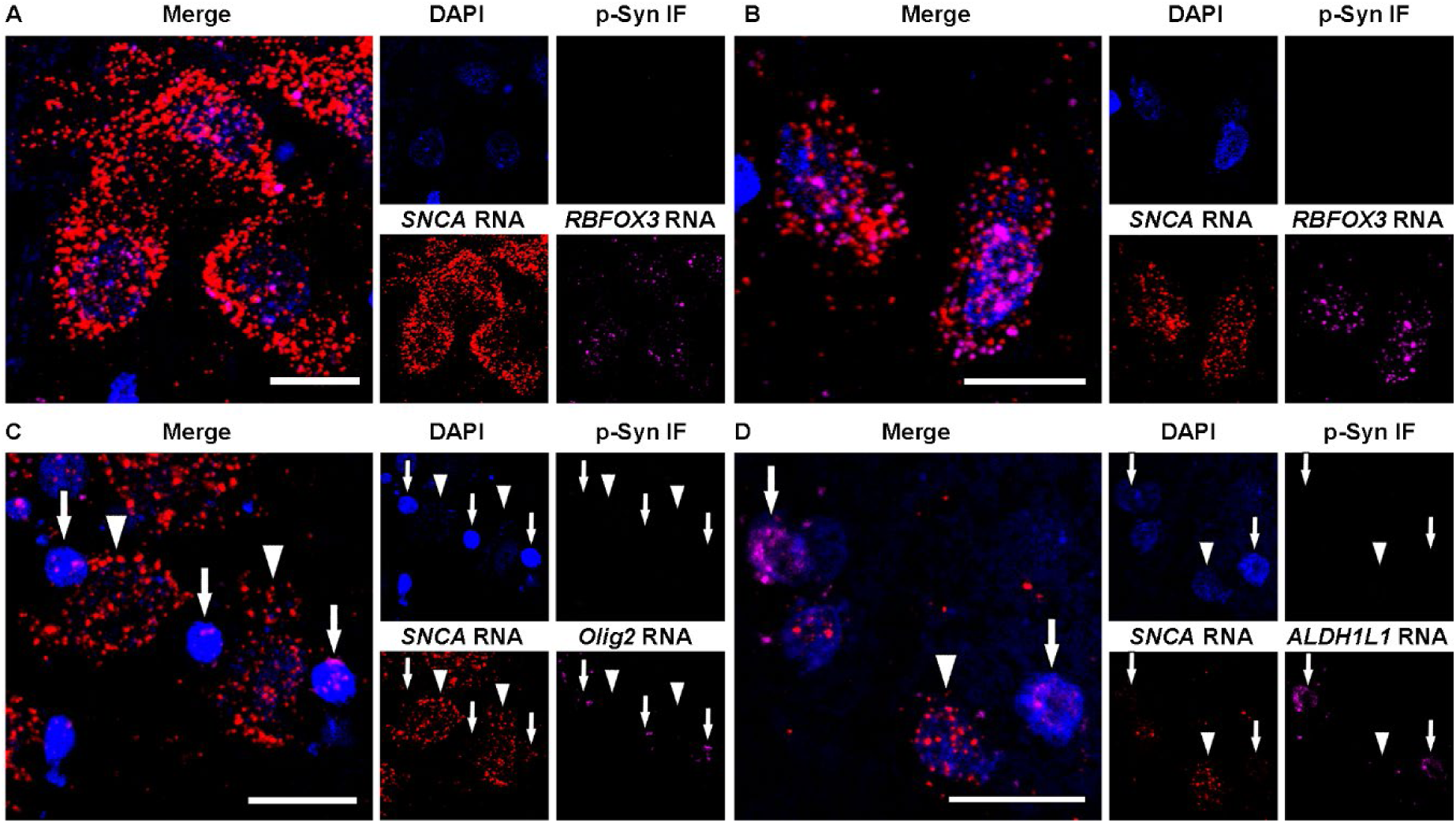
Representative images of RNAscope combined with immunofluorescence in control cases. **A, B** Neurons in the substantia nigra (**A**) and amygdala (**B**) express numerous *SNCA* and neuron-specific *RBFOX3* transcripts in the nucleus and cytoplasm. **C** Oligodendrocytes contain many *Olig2* transcripts and few *SNCA* transcripts (arrows) whereas neurons exhibit numerous *SNCA* transcripts and an absence of *Olig2* transcripts (arrowheads). **D** Astrocytes contain many *ALDH1L1* transcripts and few *SNCA* transcripts (arrows) while neurons show numerous *SNCA* transcripts and an absence of *ALDH1L1* transcripts (arrowhead). Each image set represents confocal images taken from the same field of view showing DAPI (blue), phosphorylated-α-syn (p-Syn) immunofluorescence (green), *SNCA* (red), neuron-specific *RBFOX3* (**A, B**), oligodendrocytes-specific *Olig2* (**C**), or astrocyte-specific *ALDH1L1* (**D**, magenta), and the enlarged merged image. Note that in control cases p-Syn pathology is not detected. Scale bars represent 20 μm.

**Fig. 2.**
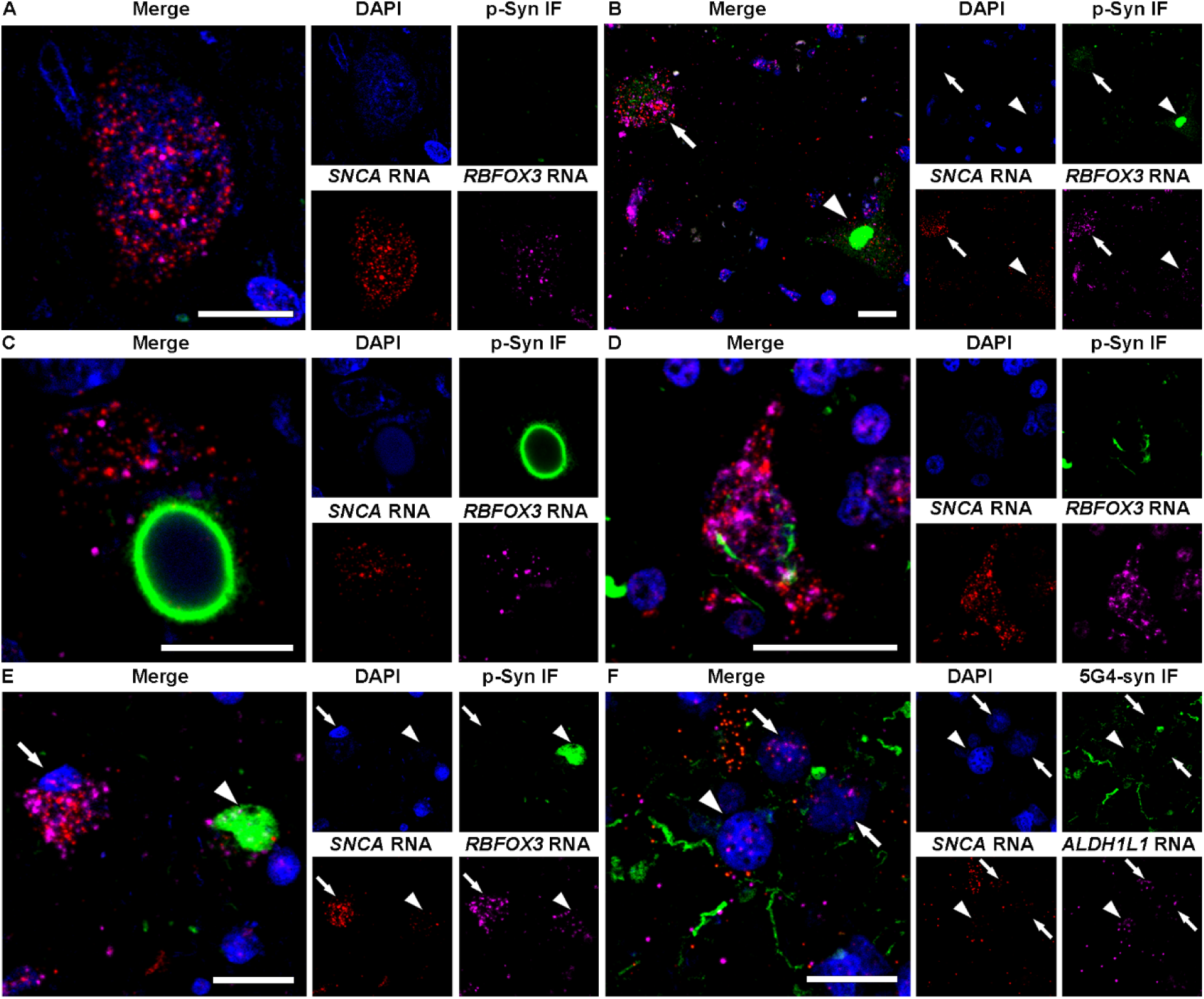
Representative images of RNAscope combined with immunofluorescence in Lewy body disease cases. **A** A neuron without α-synuclein (α-syn) immunoreactivity (IR) in the substantia nigra, which has numerous *SNCA* and neuron-specific *RBFOX3* transcripts in the nucleus and cytoplasm. **B** Punctate α-syn IR (arrow) in a neuron, which shows many fine dots-like α-syn IR, and an irregular-shaped compact inclusion (arrowhead), which shows compact α-syn IR without a core or halo. Numerous *SNCA* transcripts are observed in the neuron containing the punctate α-syn IR compared with the neuron with irregular-shaped compact inclusion. **C** Brainstem-type Lewy body (bLB) containing a round core and pale halo shows few *SNCA* transcripts. **D**. A neuron containing a punctate α-syn IR in the amygdala shows numerous *SNCA* transcripts. **E**. A neuron without α-syn IR (arrow) and a cortical LB (cLB, arrowhead) in the amygdala show few *SNCA* transcripts. **F**. Astrocytes with (arrowhead) and without (arrows) α-syn IR in the amygdala contain few *SNCA* transcripts. Each image set represents confocal images taken from the same field of view showing phosphorylated-α-syn (p-Syn, **A-E**) or 5G4 α-syn (5G4-syn, **F**) immunostaining (green), *SNCA* (red), neuron-specific *RBFOX3* (**A-E)** or astrocyte-specific *ALDH1L1* (**F**, magenta), DAPI (blue), and the enlarged merged image. Scale bars represent 20 μm.

Pathological α-syn immunoreactive neurons were stratified as those with more than two small dot-like or threadlike immunostaining (so-called punctate α-syn IR) [8-10], and those with compact inclusions. Compact inclusions in the substantia nigra were further subclassified as irregular-shaped compact inclusions which did not correspond to the classical brainstem-type LB (bLB), and the classical bLB with a round core and pale halo [8-10]. In the amygdala, we distinguished punctate α-syn IR and compact inclusions identified as cortical LB (cLB) [10]. To quantify the nuclear area density of *SNCA* transcripts within neurons, astrocytes, and oligodendrocytes, an ROI encompassing the nucleus, visualized using DAPI staining, was meticulously delineated. This analysis was performed on neurons in the SN and amygdala, oligodendrocytes in the pons, and astrocytes in the amygdala using 2 control cases.

### snRNA-seq and data processing

snRNA-seq was performed as previously reported [38]. In brief, 3 control cases of frozen frontal cortex stored at -80℃ were chopped and loaded on Singulator TM 100 (S2 Genomics1). Small-volume nuclei isolation was performed using the Automated Tissue Dissociation System with minor adjustments. Singulator cartridges (S2 Genomics) were inserted with frozen tissues and mounted on the system for nuclei isolation. The collected nuclei were stained with DAPI and sorted using an Aria Fusion A cell sorter for DAPI-positive cells. The nuclei were stained with SYBR Green II and counted under a microscope using INCYTO C-Chip hemocytometers (Neubauer Improved). Sorted nuclei were used as input into the 10X Genomics single-cell 3’ v3.1 assay and processed as 10X Genomics protocol and then the library was constructed. The quality of the library was assessed by Bioanalyzer (Agilent Technologies) and qPCR amplification data (Roche).

The single-nuclei transcriptome sequencing data were processed using the Cell Ranger Single-Cell Software Suite (v 6.0.0) from 10X Genomics. The raw data were demultiplexed to identify cell and unique molecular identifier (UMI) barcodes, followed by alignment to the GRCh38 reference genome using the STAR (v 2.7.10) tool available in the Cell Ranger pipeline. Gene expression quantification was combined into a single feature-barcode matrix. To normalize the depth across all merged datasets, the ’Cell Ranger aggr’ function was utilized, ensuring a similar number of uniquely mapped transcriptome reads per cell.

Dimensionality reduction was performed using principal component analysis (PCA), with the top 10 principal components selected for subsequent analyses. Cells were clustered based on the k-means algorithm, and the results were visualized using UMAP (Uniform Manifold Approximation and Projection). Differentially expressed genes (DEGs) were identified using the Seurat FindConservedMarkers function (p <0.05; negative binomial exact test). To assign cell types, known brain cell type markers [40] were mapped to the DEGs in each cluster. Three approaches were employed to assign cell type identity with stringent criteria. Initially, we created a graph showing the count of shared genes between known marker genes and cluster-specific top 50 differentially expressed genes in each cluster. Second, we generated a plot illustrating the average expression levels of marker genes that were differentially expressed across all clusters. Third, we calculated the average expression levels of the known marker genes in various clusters. Cell type identity was assigned to each cluster based on the restricted expression of marker genes, incorporating information from all three approaches. Heatmaps and boxplots were generated using the plotly package in Python.

### Statistical analysis

Statistics were performed using GraphPad Prism (version 9) and SPSS Statistics Version 23. Kruskal-Wallis test and Dunn’s post hoc analysis with Bonferroni correction were used to compare the area density of *SNCA* transcripts. Mann Whitney test with Bonferroni correction were used to compare the expression of *SNCA* using snRNA-seq data. The 10^th^ percentile Z-score cut-off was calculated by Statology (https://www.statology.org/z-score-cut-off-calculator/) to determine the proportion of the area density of LBs below the 10^th^ percentile of the neurons without α-syn IR [41]. If this proportion reached 50% of cells, we interpret this as a strong indicator for a decrease in SNCA transcript area density compared with neurons without α-syn IR [38]. Two-sided p <0.05 was considered significant.

### Ethical considerations

This study was approved by the University Health Network (UHN) Research Ethics Board (Nr. 20-5258) and the University of Toronto (Nr. 39459) and was performed per ethical standards established in the 1964 Declaration of Helsinki, updated in 2008.

## Results

### Demographics and neuropathological data of cases

Demographics and neuropathological data of patients used in this study are summarized in **Table 1**. Age in LBD cases ranged from 62 to 85 (median 73) years at death, and control cases ranged from 52 to 77 (median 70) years at death. Postmortem interval in LBD cases ranged from 4 to 24 (median 15) hours, and control cases ranged from 4 to 30 (median 12.5) hours. Braak LBD stages in LBD were 4 and 5, and no Lewy pathology was observed in control cases. ADNC was intermediate in 2 cases of LBD, and absent in other cases.

### RNAscope combined with immunostaining for α-syn

In the control cases (n = 2), *SNCA* transcripts were observed abundantly in the nucleus and cytoplasm in neurons in the SN, amygdala, and pons (**Fig. 1A-C**), while only single *SNCA* transcripts were found in the nucleus and cytoplasm in the oligodendrocytes in the pons (**Fig. 1C**) and in astrocytes in the amygdala (**Fig. 1D**). *SNCA* transcripts were clearly distinguishable from autofluorescence neuromelanin or lipofuscin particles (**Fig. S1**). In accordance with our previous report [38], transcripts of *RBFOX3*, *Olig2*, and *ALDH1L1* in neurons, oligodendrocytes, and astrocytes, respectively, were detected both in the cytoplasm and nucleus in each cell type. Phosphorylated α-syn IR was not observed in the control cases.

In LBD cases, *SNCA* transcripts were also observed in the neuronal nucleus and cytoplasm (**Fig. 2A, E**) including those containing punctate α-syn IR (**Figure 2B, D**), irregular-shaped compact inclusion (**Fig. 2B**), bLB (**Fig. 2C**), and cLB (**Fig. 2E**). However, *SNCA* transcripts were only rarely found in the α-syn immunoreactive LB areas (**Fig. 2C, E, Additional file 2**). As reported previously, our investigations employing phosphorylated α- syn antibody failed to identify α-syn IR in astrocytes. Instead, we utilized the 5G4 α-syn antibody, known for its ability to detect disease-associated astrocytic α-syn IR [21, 39, 42, 43]. However, *SNCA* transcripts were infrequently observed both with and without disease-associated α-syn IR in astrocytes (**Fig. 2F**).

### Quantification of *SNCA* transcripts in various α-syn cytopathologies

Due to significant neuronal loss in LBD cases, a limited number of residual neurons (ranging from 17 to 50) were observed per section in the SN. *SNCA* area density was similar in pooled neurons without α-syn IR (n = 117 in the SN, median 17.1%; n = 305 in the amygdala, median 27.9%) and pooled neurons with punctate α-syn IR (n = 18 in the SN, median 13.8%; n = 37 in the amygdala, median 26.8%) (**Fig. 3A, B**). However, a gradual decrease of *SNCA* area density in pooled neurons with irregular-shaped compact inclusions (n = 14, median 12.4%) followed by pooled bLBs (n = 17, median 5.6%) was observed, which showed statistical significance (neurons without α-syn IR vs bLB, p <0.001; neurons with punctate α-syn IR vs bLB, p =0.01) in the SN (**Fig. 3A**). Also, the area density of *SNCA* transcripts in pooled neurons with cLBs (n = 176, median 14.1%) was significantly lower than those without α- syn IR (p <0.001) or with punctate α-syn IR (p <0.001) in the amygdala (**Fig. 3B**). Distribution of area density of *SNCA* transcripts in each morphological group was displayed in the **Fig. 3C and D**. Although *SNCA* transcript area density varied between pooled neurons with and without α-syn IR, the distribution curves gradually deviated to the lower values during the maturation process of LB formation (**Fig. 3C, D**). Among the observed inclusions, 14.3% of irregular-shaped compact inclusions, 35.3% of bLBs, and 69.3% of cLBs fell below the 10th percentile Z-score cut-off of neurons without α-syn IR.

**Fig. 3.**
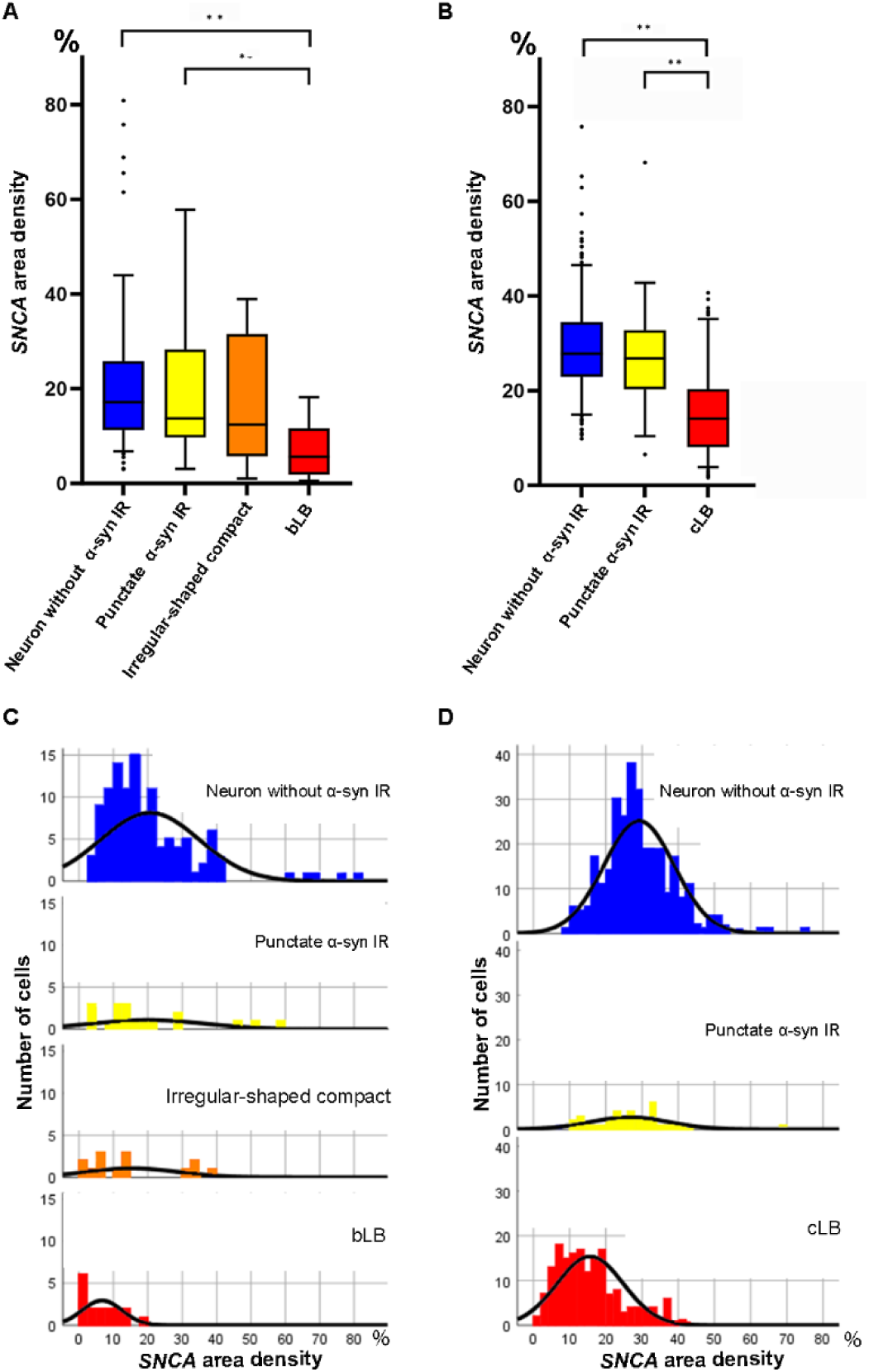
Area density of *SNCA* transcripts in neurons in the substantia nigra and amygdala. **A, B** Box and whisker plots of *SNCA* transcripts area density in the substantia nigra (SN, **A**) and amygdala (**B**). *SNCA* transcript area density in neurons without α-synuclein (α-syn) immunoreactivity (IR) in the SN (n = 117) and amygdala (n = 305) are similar to neurons with α-syn IR in the SN (n = 18) and amygdala (n = 37). However, *SNCA* transcript area density in neurons with irregular-shaped compact inclusions (n = 14), and those with brainstem-type Lewy bodies (bLB, n = 17) in the SN show a gradual decrease (**A**). The *SNCA* transcript area density in neurons containing cortical LB (cLB, n = 176) also shows a decrease compared to neurons without α-syn IR and punctate α-syn IR (**B**). The upper and lower whiskers indicate the 95th percentile and 5th percentile, respectively. The top and bottom of the box indicate the first and third quartiles, respectively. The middle line splitting the box indicates the median. The small circles in the plots indicate outliers. For statistics, Kruskal-Wallis test and Dunn’s post hoc analysis with Bonferroni correction are used. Statistically significant findings where p <0.05 and p <0.001 are indicated as * and **, respectively. **C, D** Distribution of *SNCA* area density in neurons in the SN (**C**) and amygdala (**D**). Distribution curves of *SNCA* area density gradually deviate to a lower value during the maturation process of LBs. X-axis represents the area density expressed as the percentage and y-axis demonstrates the number of cells (Note that y-axis scales are different from **C** and **D**).

### Nuclear *SNCA* transcripts area density

We further quantified *SNCA* area density within the nucleus in neurons, oligodendrocytes, and astrocytes in control cases (n = 2) to compare with the results of snRNA-seq. We found that the nuclear area density of *SNCA* transcripts in pooled neurons (n = 101 in the SN, median 30.1%; n = 112 in the amygdala, median 37%) was significantly higher (p <0.001) compared to pooled oligodendrocytes (n = 113, median 8.6%) and pooled astrocytes (n = 105, median 5.3%). Furthermore, the nuclear area density of *SNCA* transcripts in oligodendrocytes was significantly higher than that in astrocytes (p <0.05). No statistically significant distinction in the nuclear *SNCA* transcripts area density was observed between neurons in the SN and amygdala (**Fig. 4**).

**Fig. 4.**
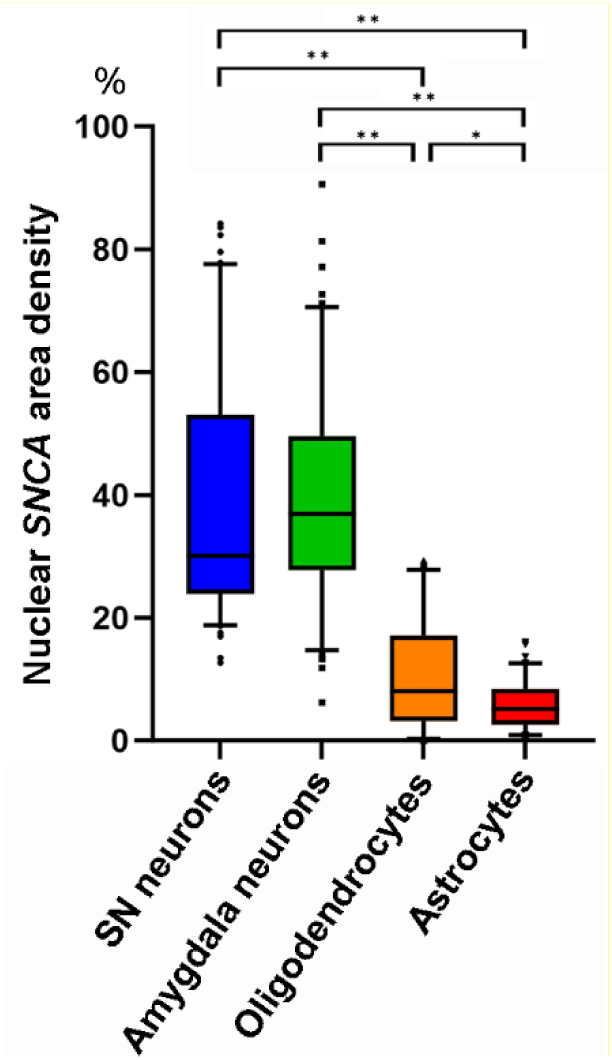
Nuclear area density of *SNCA* transcripts in neurons, oligodendrocytes, and astrocytes in controls. Box and whisker plots of nuclear area density of *SNCA* transcript area density in neurons in the substantia nigra (SN, n = 101) and amygdala (n =112), oligodendrocytes in the pons (n = 113), and astrocytes in the amygdala (n = 105) in control cases. The nuclear area density of *SNCA* transcripts in neurons is significantly higher than those in oligodendrocytes and astrocytes (p <0.001). The nuclear area density of *SNCA* transcripts in oligodendrocytes is significantly higher than that of astrocytes (p <0.05). The upper and lower whiskers indicate the 95th percentile and 5th percentile, respectively. The top and bottom of the box indicate the first and third quartiles, respectively. The middle line splitting the box indicates the median. The small circles in the plots indicate outliers. For statistics, Kruskal-Wallis test and Dunn’s post hoc analysis with Bonferroni correction are used. Statistically significant findings where p <0.05 and p <0.001 are indicated as * and **, respectively.

### Cell-type specific *SNCA* expression in the frontal cortex

Based on our RNAscope analysis, we detected occasional *SNCA* transcripts in oligodendrocytes and astrocytes. This observation would add to the current incomplete understanding of α-syn expression in glial cells because α-syn is generally considered a neuronal protein [44, 45]. Therefore, we further conducted snRNA-seq analysis in control cases (n =3). A comprehensive dataset comprising 15,265 transcriptome profiles from individual nuclei was successfully generated. The analysis revealed a median count of 8,007 unique molecular identifiers (UMIs) per cell, corresponding to a median detection of 2,998 genes per cell. We achieved an average sequencing saturation of 88.8% for the libraries. Following this, we performed cluster annotation based on the expression of established marker genes associated with known cell types. Our snRNA-seq analysis uncovered a distinct pattern specific to each cell type (**Fig. 5A**). Notably, we observed a substantial expression of *SNCA* transcripts in homeostatic microglia, oligodendrocyte progenitor cells, and inhibitory neurons (p = 4.4×10^-5^ vs mature oligodendrocytes, p = 4.8×10^-26^ vs astrocytes). Conversely, we noted a low expression in excitatory neurons (p = 7.3×10^-21^ vs astrocytes) and mature oligodendrocytes (p = vs 1.2×10^-33^ vs astrocytes). Moreover, the expression of *SNCA* transcripts was largely absent in astrocytes, microglia, and endothelial cells (**Fig. 5B**). It is noteworthy that the expression of the *SNCA* transcript is significantly higher in inhibitory neurons compared to excitatory neurons (p = 3.3×10^-6^).

**Fig. 5.**
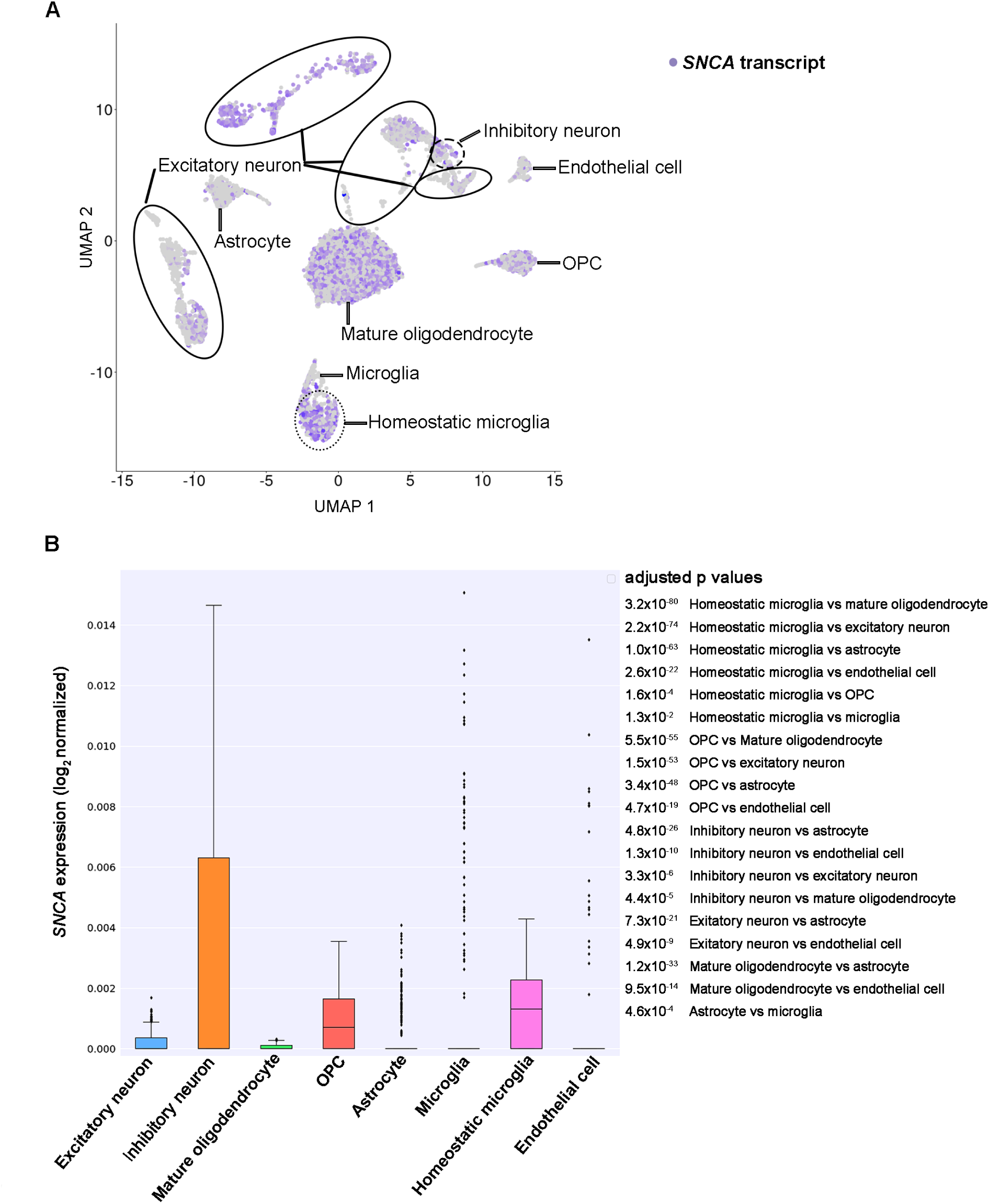
*SNCA* expression in single-nucleus RNA sequencing in control cases. **A** Expression profile of *SNCA* in major cell types of the control frontal cortex uncovering a distinct pattern specific to each cell type. Each point represents a single cell and color intensity indicates the normalized expression level of *SNCA*. UMAP projection of 15, 265 brain cells included. **B** Quantification of *SNCA* expression across cell types. The median of expression is depicted as a line within the box and whiskers indicate the range of the normalized *SNCA* expression. For statistics, Mann Whitney test with Bonferroni correction are used. Adjusted p values are listed on the right side of the figure. OPC, oligodendrocyte progenitor cell.

### Immunohistochemistry for pan-and disease-associated-α-syn antibodies

To support literature data on preserved α-syn protein expression in LBD, we immunostained SN and striatum using an antibody that detects the physiological and disease-associated form of α-syn and one that detects the disease-associated α-syn. This confirmed preserved synaptic immunoreactivity in the striatum and SN for the antibody detecting the physiological form of α-syn in both non-diseased controls and LBD samples, while only LBD samples showed typical disease-associated α-syn deposits using the 5G4 antibody. For details see **supplementary results** and **Fig. S2**.

## Discussion

This study revealed preserved, but gradually decreasing, *SNCA* transcription in the SN and amygdala neurons during the maturation process of LB formation.

‘Proteinopathy’ and ‘proteinopenia’, representing contrasting concepts, dictate divergent treatment strategies: the elimination of disease-associated α-syn or the replacement of normal physiological α-syn. *SNCA* transcripts in the SN in LBD have been reported to be both up-and down-regulated compared with controls [26-35]. The previous studies used a tissue-digested method that reported the average expression levels across a complex of neuronal and non-neuronal cell types. Increased *SNCA* transcripts in neuromelanin-containing neurons in the substantia nigra in Parkinson’s disease compared with controls were reported using laser-microdissection and quantitative reverse transcription PCR [27].

In contrast to our cytopathology-based evaluation, the approach of that study was not able to compare *SNCA* transcripts and α-syn pathology. The individual dots of RNAscope signal represents its expression level of transcripts [46]. However, *SNCA* transcripts appeared as isolated dots or in small confluent clusters, making it difficult to accurately count individual transcripts. Therefore, we used the area density measurements of *SNCA* transcripts as its expressional level in this study that reliably detects alterations in the transcripts [38].

We noted a variation in *SNCA* transcripts within morphologically distinct α-syn IR cytopathologies (**Fig. 3C, D**), with the mature LBs showing the least. This diversity in *SNCA* expression likely reflects a dynamic phenomenon in the human brain, as transcription at the cellular level undergoes continuous fluctuations in response to physiological or pathological demands. Similarly, protein expression of mitochondrial complex markers has been reported to differ between α-syn cytopathologies, supporting the concept that α-syn deposition is associated with dynamic cellular protein and RNA responses [47, 48]. While the relationship between *SNCA* mRNA and protein expression is complex [49, 50], proteins are synthesized from mRNA templates [51]. Therefore, it is commonly observed that transcript expression levels correlate with protein synthesis [16, 17, 19, 51, 52]. Interestingly, for another neurodegenerative disease-related protein, tau, the regional variability in total tau protein expression levels correlates with similar changes in mRNA expression levels evaluated with the quantitative RT-PCR method [52]. Indeed, and in line with previous studies using Western blots [6, 39, 53, 54], demonstrating preserved expression of monomeric α-syn in LBD samples, we also demonstrate preserved monomeric α-syn IR in the SN and striatum of LBD cases (**Fig. S2**).

Our RNAscope analysis revealed a decrease in *SNCA* transcripts during the maturation process of LB formation. Most likely this reflects an exhaustion of the transcription due to the constant production of α-syn that is then utilized for the seeding of misfolded α-syn. This phenomenon would be reminiscent of the mechanism described in the leading model of neurodegenerative proteinopathies, which is associated with misfolded prion protein (PrP), wherein the progressive accumulation of misfolded PrP leads to a depletion of the pool of physiological cellular PrP [55]. Considering *SNCA* transcripts are rarely observed within the region of α-syn immunoreactive LB as observed through RNAscope, it can also be hypothesized that neurons are unable to generate *SNCA* transcripts due to the densely packed filaments that constitute the LBs’ ultrastructural composition [9, 56, 57]. Finally, we cannot exclude the possibility that the accumulation of misfolded α-syn during LB maturation exerts a negative feedback effect on the transcriptional regulation of *SNCA*.

Based on our observations we hypothesize the following scenario (**Fig. 6**). Neurons have a well-described maturation process of α-syn protein IR [8-10], driven also by posttranslational modifications affecting the amplification of disease-associated α-syn [58], which include punctate α-syn IR, irregular-shaped compact inclusion, and classical LBs [2, 8, 9, 11, 12, 15]. Neurons contain *SNCA* transcripts to serve as a pool for protein production. Following a “neurodegenerative event” (it is unclear what triggers the process; this may vary considerably from patient to patient), monomeric α-syn misfolds, and serves as seeds for further disease propagation [1, 2, 4, 5]. In addition, α-syn seeds are internalized from the extracellular space [2, 21]. Morphologically, an early cytopathological alteration reflecting this process is the punctate non-ubiquitin positive α-syn IR. The neuron recognizes the misfolded α-syn and attempts to ‘sequester’ it leading to the formation of a LB. Due to the non-physiological misfolded α-syn, the neuron attempts to preserve physiological functioning by maintaining normal levels of *SNCA* transcripts. At a certain stage in the maturation process of protein aggregation, neurons reach a critical state and either 1) exhaust their cellular pool of *SNCA* transcripts, resulting in reduced *SNCA* transcripts and in parallel, segregate the misfolded protein into a LB or, 2) can no longer process the misfolded α-syn and degenerate leading to neuronal loss and detection of extracellular neuromelanin in neuropathology. Importantly, the *SNCA* transcription and physiological monomeric α-syn expression are not lost completely, as shown by the preserved physiological presynaptic α- syn staining and Western blot observations [6, 39, 53, 54]. Regarding astrocytes, the lack of cellular production of *SNCA* strongly supports the notion that astrocytes ingest disease-associated α-syn [21]. This contrasts the development of astrocytic tau pathology where astrocytes can produce tau themselves to serve as a local pool for misfolding [38].

**Fig. 6.**
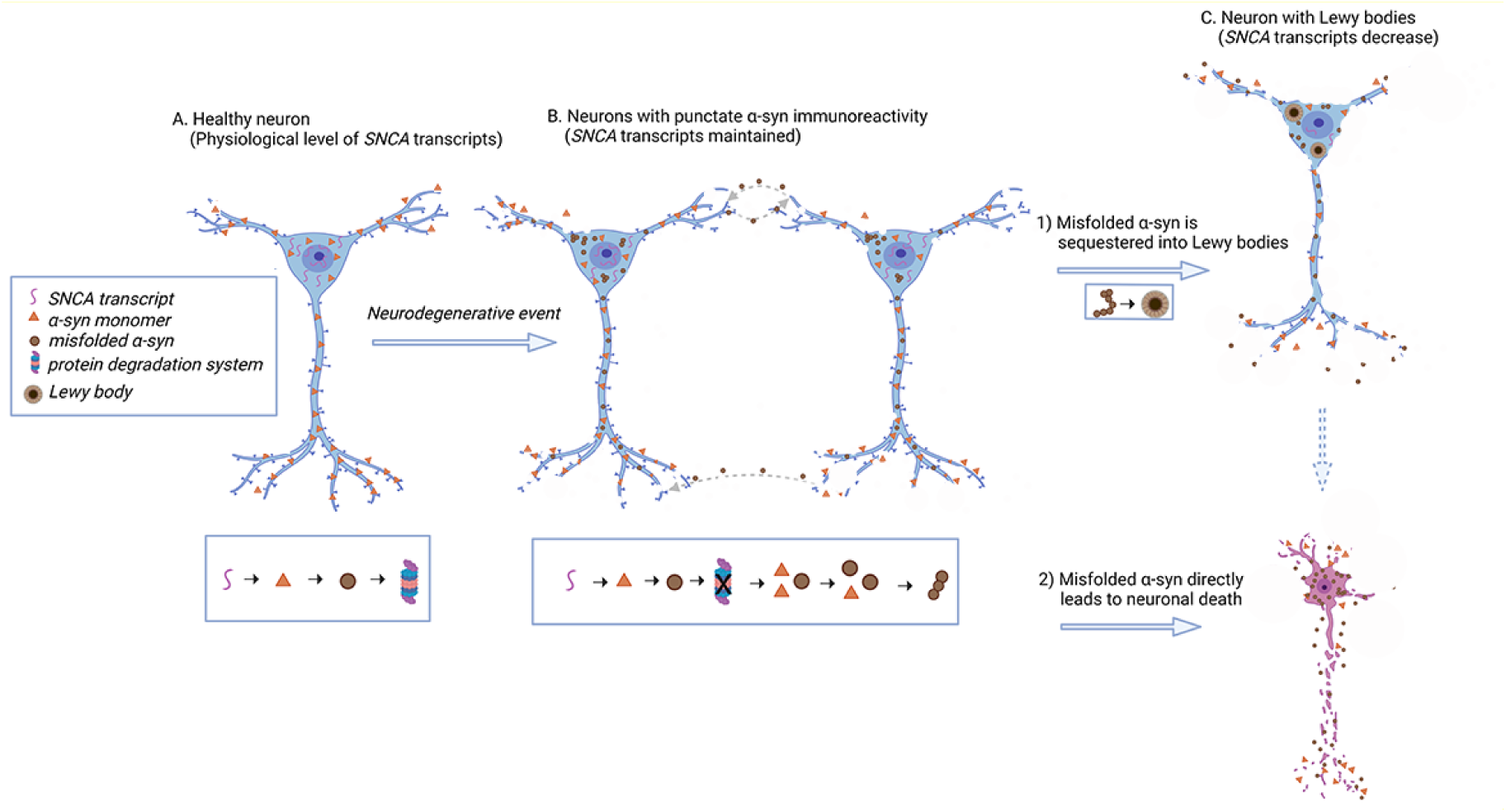
Schematic hypothesis of cellular regulation of *SNCA* transcripts during the maturation process of Lewy body formation. **A** In healthy condition, neurons express physiological levels of *SNCA* transcripts, and their protein degradation systems function normally. **B** When a “neurodegenerative event” occurs, (e.g. protein degradation system dysfunction) monomeric α-synuclein (α-syn) misfolds and serves as seeds for further disease propagation. Additionally, α-syn seeds are internalized from the extracellular space. Punctate α-syn immunoreactivity is observed in the cytoplasm in both cases. The neuron tries to ‘sequester’ misfolded α-syn. Due to the misfolding of α- syn, its physiological function is impaired. The neuron attempts to preserve physiological functioning by maintaining normal levels of *SNCA* transcripts. **C** Neurons reach a critical state and either 1) the misfolded α-syn is segregated into Lewy body, *SNCA* transcripts reduce due to cell dysfunction, ultimately leading to neuronal death, or, 2) the misfolded α-syn directly leads to neuronal death. Expression level of monomeric α-syn is preserved during the progression of the disease. Created with BioRender.com

We infrequently observed *SNCA* transcripts in oligodendrocytes and astrocytes by RNAscope analysis. Nuclear *SNCA* area density in these glial cells was significantly lower than that of neurons, in addition, that of oligodendrocytes was significantly higher than astrocytes. Our snRNA-seq findings support the results, showing high *SNCA* transcripts expression in inhibitory neurons, OPCs, and homeostatic microglia, while low expression in excitatory neurons and mature oligodendrocytes. In contrast, *SNCA* transcripts were hardly detectable in astrocytes, microglia, and epithelial cells. Notably, SNCA transcript expression in inhibitory neurons was significantly higher than in excitatory neurons. In mice cultured cells, α-syn aggregate formation was observed in GAD-positive inhibitory neurons [59, 60]. Furthermore, α-syn positive inclusions were identified in medium spiny neurons, known for GABAergic inhibitory cells, in patients with PD [61]. Given that α-syn expression levels are directly associated with LB pathology [62], the enrichment of *SNCA* transcripts in specific neuronal cell types may influence their vulnerability to LB pathology. Oligodendrocytes, astrocytes, and microglia appear to express α-syn, but the level of expression is much lower than that in neurons [63-67]. We compared our *SNCA* transcripts expression dataset in snRNA-seq with 3 publicly available databases: 1) a human single-cell RNA-seq database, scRNAseqDB (https://bioinfo.uth.edu/scrnaseqdb/), 2) single cell RNAseq section in THE HUMAN PROTEIN ATLAS (HPA, https://www.proteinatlas.org/), 3) an immunopanning purification cell based RNA-Seq database, Brain RNA-Seq database (http://www.brainrnaseq.org, refers to an article [68]). Consistent with our dataset, we observed moderate to high *SNCA* transcript expression in neurons and OPCs, and low to moderate expression of *SNCA* transcripts in oligodendrocytes, while astrocytes exhibited low expression across all databases. However, there were variations in the expression levels of *SNCA* transcripts in microglia among the different databases. Our data exhibited high expression of *SNCA* transcripts in homeostatic microglia, while low expression in microglia. Brain RNA-Seq showed moderate *SNCA* transcript expression in microglia, whereas both scRNAseqDB and HPA indicated low expression. *SNCA* transcript expression in homeostatic microglia was not accessible in these databases. Mice with *SNCA* overexpression exhibit microglial activation, which contributes to the degeneration of dopaminergic neurons [69]. The expression of *SNCA* transcripts may play a role in microglial cell activation/differentiation and contribute to neurodegenerative processes. Notably, our data was obtained through single ’nucleus’ RNA-seq, whereas the other databases utilized single ’cell’ RNA-seq methodologies. Therefore, our study highlights nuclear *SNCA* transcripts expression can be different from cytoplasm or whole-cell transcripts. By employing snRNA-seq, it has been revealed that *SNCA* is not limited to dopaminergic neurons but is also expressed in other neurons, oligodendrocytes, and homeostatic microglia. This observation challenges the conventional notion of α-syn as exclusively a neuronal protein [3], shedding light on its expression in glial and microglial cells. The significance of *SNCA* transcript expression underlying the formation of oligodendroglial cytoplasmic inclusions, which serve as a distinctive pathological feature of multiple system atrophy (MSA), is emphasized. We highlight our ongoing efforts to investigate the expression of *SNCA* in oligodendroglial cytoplasmic inclusions in MSA, aiming to provide further insights into the disease.

It is important to note that this study has some limitations. Detailed morphometric evaluations can be performed on only a small number of cases. However, the cytopathologies were well represented and the preserved *SNCA* transcripts were clear in all. In addition, we did not assess the expression of *SNCA* transcripts in specific cell lineages, such as dopaminergic neurons, GABAergic neurons, and immature astrocytes. This is because the RNAscope method used here does not allow the application of a wide range of neuronal markers. Importantly, we focused on neuromelanin-containing cells of the substantia nigra that are dopaminergic.

## Conclusions

Our study revealed preserved but gradually decreasing *SNCA* transcription in substantia nigra and amygdala neurons during the maturation process of LB formation. Our study advances our understanding of the pathogenesis of LBD by elucidating the upstream regulation of *SNCA* expression during disease progression and uncovering cell-type specific diversity of *SNCA* expression. These findings provide valuable insights into the molecular mechanisms underlying pathogenesis and offer guidance for a therapeutic approach to disease modification in LBD. Importantly, our study provides a rationale for a cautiously balanced dual approach to therapies obstructing α-syn aggregation and in parallel reducing *SNCA* transcription to avoid feeding the misfolding process by the production of α-syn. We highlight novel aspects of disease pathogenesis that will be relevant for basic researchers working on cellular mechanisms of synucleinopathies.

## Abbreviations

α-syn: α-Synuclein; ACD: Advanced cell diagnostics; ADNC: Alzheimer’s disease neuropathological change; ALDH1L1: Aldehyde dehydrogenase 1 family member L; bLB: Brainstem-type Lewy body; cLB: Cortical Lewy body; DAPI: 4′,6-diamidino-2- phenylindole; DEG: Differentially expressed gene; HPA: HUMAN PROTEIN ATLAS; IHC: Immunohistochemistry; IR: Immunoreactivity; LB: Lewy body; LBD: Lewy body disease; MSA: Multiple system atrophy; Olig2: Oligodendrocyte transcription factor 2; OPCs: Oligodendrocyte progenitor cells; PCA: Principal component analysis; PCR: Polymerase chain reaction; RBFOX3: RNA Binding Fox-1 Homolog 3; ROI: Region of interest; SN: Substantia nigra; SNCA: α-Synuclein gene; snRNA-seq: Single nucleus RNA sequencing; UMAP: Uniform manifold approximation and projection; UMI: Unique molecular identifier

## Acknowledgments

The authors particularly acknowledge the patients and their families for their donations. Fig. 6 was designed with BioRender.com.

## Authors’ contributions

TK carried out the RNAscope and IHC experiments, analyzed and interpreted the data, and drafted the manuscript. SLF conducted the IHC experiment, analyzed and interpreted the data, and edited the manuscript. SL performed the RNAscope and snRNA-seq experiments, analyzed and interpreted the data, and edited the manuscript. IMV carried out the snRNA-seq experiment, interpreted the data, and edited the manuscript. JL conducted the RNAscope and IHC experiments and edited the manuscript. NN performed the snRNA-seq experiment and analyzed and interpreted the data, and edited the manuscript. MJU carried out the snRNA-seq experiment, analyzed and interpreted the data, and edited the manuscript. AEL interpreted the data and edited the manuscript. GGK conducted the RNAscope and IHC experiments, analyzed and interpreted the data, and edited the manuscript. All authors read and approved the final manuscript.

## Availability of data and materials

All data generated and analyzed during the current study are included in this article and are available from the corresponding author upon reasonable request.

## Consent for publication

Not applicable.

## Supplementary information

### Supplementary methods

#### Immunohistochemistry

For immunohistochemistry, midbrain and basal ganglia (n = 5 each in cases of LBD and controls) were analyzed. To assess the presence of physiological monomeric α-syn protein, we utilized immunohistochemistry with two α-syn specific antibodies: SYN-1(1:2000, BD Biosciences, Franklin Lakes, NJ) [1], which is known to cross-react with physiological monomeric α-syn and disease-associated one [1], and 5G4, which specifically targets disease-associated α-syn protein and does not exhibit IR against physiological monomeric α-syn protein [1-3]. Antigen retrieval was performed using Dako PT Link with low pH solution and 80% formic acid for 5 minutes for 5G4 anti-α-syn antibody. According to the manufacturer’s protocol, immunostaining was performed using Dako Autostainer Link 48 and EnVision FLEX+ Visualization System. Subsequently, all sections were counterstained with hematoxylin.

**Fig. S1.**
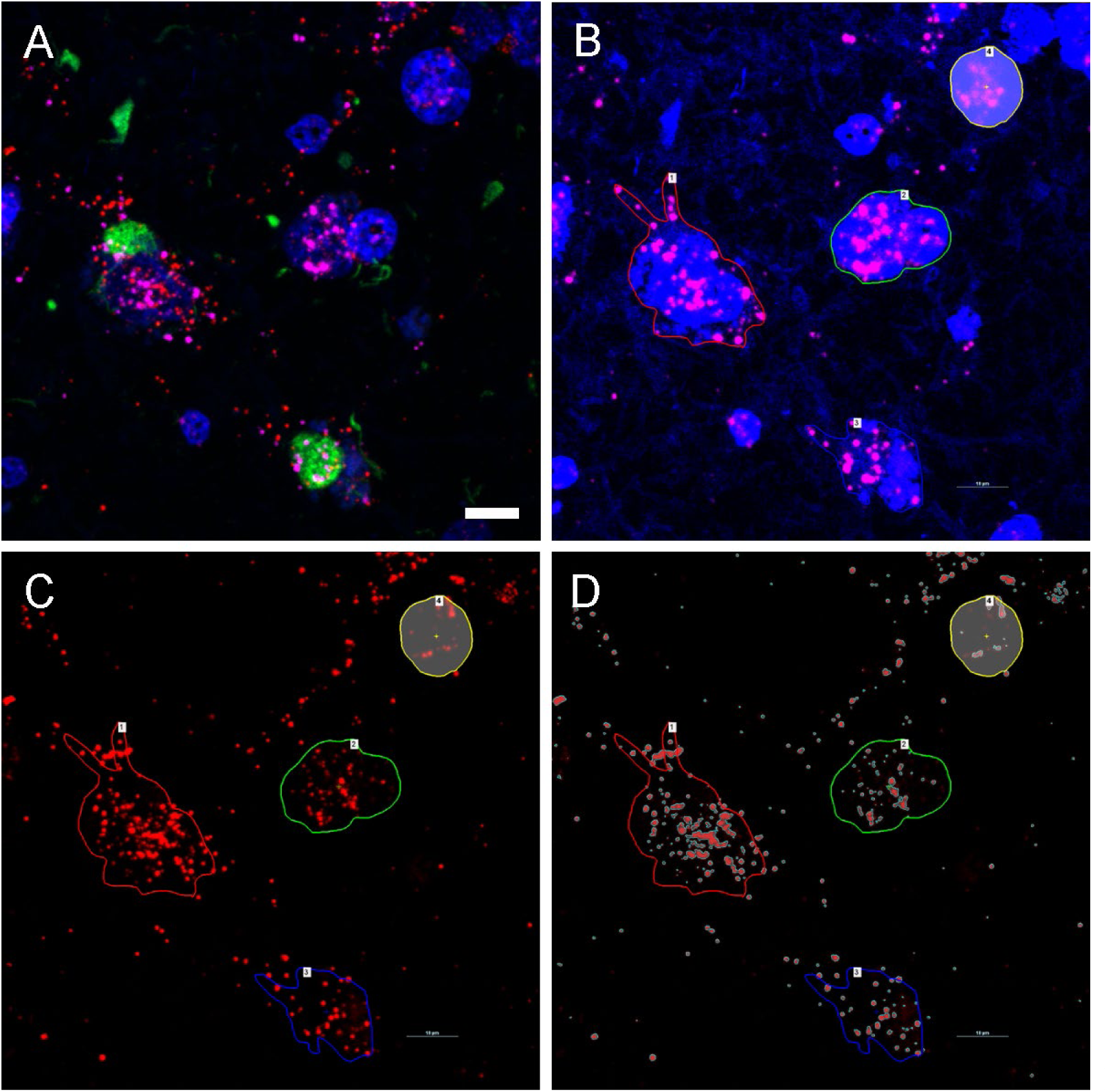
Method of outlining the region of interest and capturing the positive signals in RNAscope. Raw data image of the amygdala section in a case of Lewy body disease (**A**). The cell-specific marker and DAPI channels are selected and merged (**B**). Next, their Lookup Table strength is increased enough to make the level of autofluorescence of neuromelanin and/or lipofuscin pigments visible (**B**). Then, the cell body borders are traced by their signals (**B**). Subsequently, returning the LUT parameters to the default settings, the *SNCA* transcript channel is selected (**C**). Positive signals corresponding to *SNCA* transcripts above the threshold within the region of interest are captured by the NIS-Elements software (**D**). Green represents phosphorylated-α-syn immunostaining, red displays *SNCA* transcripts, magenta shows *RBFOX3* transcripts, and blue exhibits DAPI. Scale bar represents 10 μm.

#### Immunohistochemistry

To support literature data on preserved α-syn protein expression in LBD, for demonstration we immunostained SN and striatum using an antibody that detects the physiological and disease-associated form of α-syn and one that detects the disease-associated α-syn. We further evaluated the immunoreactivity of physiological monomeric α-syn protein in the SN and dorsal striatum where dopaminergic neurons project to their synaptic terminals in cases of control and LBD. In both control and LBD cases, SYN-1 IR displayed a synaptic staining pattern both in the SN and dorsal striatum, as well as in LBs (**Fig. S2A, B, E, F**). On the other hand, disease-associated α-syn IR using the 5G4 antibody was only detected in LBD cases (**Fig. S2C, D, G, H**).

**Fig. S2.**
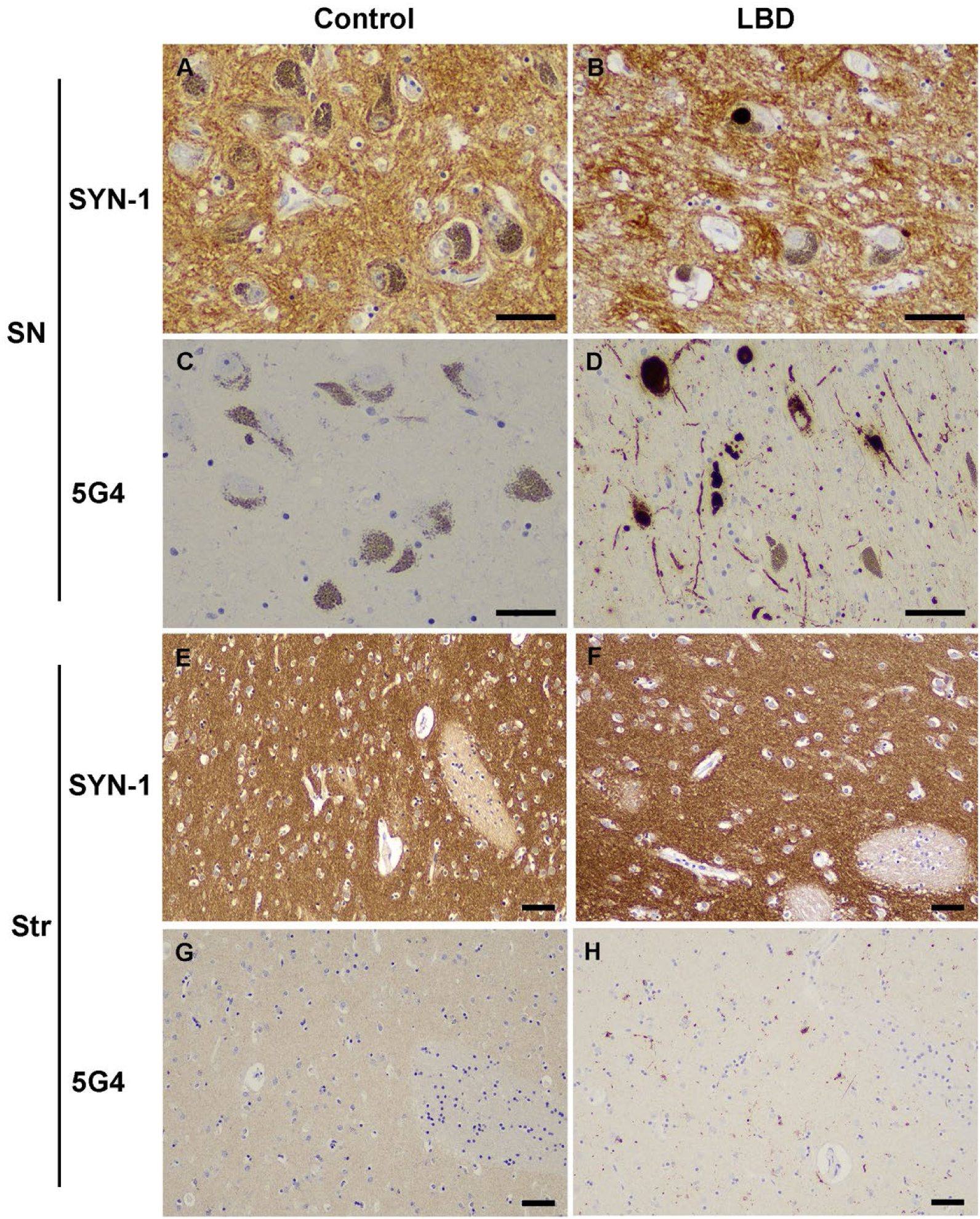
Immunohistochemistry for SYN-1 and 5G4 α-synuclein (α-syn) antibodies. SYN-1 antibody cross-reacts with the physiological monomeric α-syn and shows a synaptic pattern in both cases of controls (A, C, E, G) and LBD (B, D, F, H) in addition to revealing Lewy body and related pathology in the diseased substantia nigra (SN, D) and putamen (F). In contrast, the 5G4 antibody does not label the physiological synaptic staining in the SN and putamen in cases of control (C, G) nor of LBD (D, H) and highlights only the disease associated α-syn immunoreactivity in LBD (D, H). Immunostaining for SYN-1 (A, B, E, F) and 5G4 (C, D, G, H) anti-α-syn antibodies in the SN (A-D) and putamen (E-H). The scale bars represent 50 μm for each image.

#### Video S1. 3D RNAscope imaging of a classical Lewy body

*SNCA* transcripts are rarely observed within phosphorylated-α-syn immunoreactive Lewy body areas. Green represents phosphorylated-α-syn immunostaining, red displays *SNCA* transcripts, magenta shows *RBFOX3* transcripts, and blue exhibits DAPI. See additional video file.

